# A perceptual glitch in serial perception generating temporal distortions

**DOI:** 10.1101/2021.10.08.463190

**Authors:** Franklenin Sierra, R. Muralikrishnan, David Poeppel, Alessandro Tavano

## Abstract

Precisely estimating event timing is essential for survival, yet temporal distortions are ubiquitous in our daily sensory experience. Here, we tested whether the relative position, relative duration and relative distance in time of two sequentially-organized events —standard S, with constant duration, and comparison C, varying trial-by-trial— are causal factors in generating temporal distortions. We found that temporal distortions emerge when the first event is shorter than the second event. Importantly, a significant interaction suggests that a longer ISI helps counteracting such serial distortion effect only the constant S is in first position, but not if the unpredictable C is in first position. These results suggest the existence of a perceptual bias in perceiving ordered event durations, mechanistically contributing to distortion in time perception. We simulated our behavioral results with a Bayesian model and replicated the finding that participants disproportionately expand first-position dynamic (unpredictable) short events. Our results clarify the mechanics generating time distortions by identifying a hitherto unknown duration-dependent encoding inefficiency in human serial temporal perception, akin to a strong prior that can be overridden for highly predictable sensory events but unfolds for unpredictable ones.

## 1 Introduction

Precisely estimating event timing is essential for a range of perceptual and cognitive tasks, yet temporal distortions are ubiquitous in our daily sensory experience [1, 2, 3]. A specific kind of time distortion is the presentation-order error [4]. In 1860, Fechner observed that when comparing the weight of two elements, the order in which they were lifted mattered [5]. This led to a systematic error on subjects’ judgments of sequentially presented stimuli, which was termed time-order error (TOE) [4]. TOEs have been found in different stimulus modalities such as audition, vision and taste, as well as different stimulus dimensions such as loudness, heaviness, and brightness[4]. Understanding their mechanics is fundamental since humans normally perceive events in a series, not in isolation.

TOEs in temporal judgment can be experimentally tested by implementing a two-interval forced choice (2IFC) discrimination task, where participants compare the duration of two successive time intervals (events) per trial, a Standard and a Comparison (S vs C), separated by an inter-stimulus interval (ISI) [6, 7]. When combined with the method of constant stimuli, the duration of S remains fixed across the experimental session, whereas the duration of C changes from trial to trial and it can take one of six to nine durations distributed around the S duration [8].

In a 2IFC task, temporal performance is modelled by fitting a psychometric function. From this fitting, two main dependent variables are obtained: the point *μ* where the curve cuts the 50% line (that is, the point of subjective equality, PSE) and the slope of the resulting curve. While the PSE estimates the accuracy of the comparison judgment —and provides a marker for temporal distortions—, the slope estimates their temporal precision [9, 10]. TOE effects have been recently classified in two types: effects of the stimulus order on the PSE are called Type A effect, whereas the effects on temporal precision are called Type B effect [11, 12]. Importantly, note that there exists another type of mistake called the contraction bias: when the first stimulus is small, participants then to overestimate it, whereas when it is large, they to tend to underestimate it [13, 14].

Traditionally, TOEs have been variously attributed to sensory desensitization [15, 7], poor sensory weighting of C relative to S [16, 17], or idiosyncratic response bias [15]. More recently, two additional models have attempted to explain TOE: 1) the internal reference model (IRM) [18] —an updated version of the sensory desensitization model, and 2) Bayesian observer models [19, 20, 21].

The idea behind IRM is that in comparing S and C, participants maintain an internal representation that is the average duration of previous trials. The key idea is that this internal representation is updated by taking into consideration only the first presented duration. Because the S stimulus has a constant duration and the C stimulus varies unpredictably, more errors will be made when the order of presentation is <CS> than when it is <SC>. Raviv et al., [14] proposed a model similar to the IRM that used a Bayesian inference. Such a model assumed that the brain uses an heuristic strategy to discriminate auditory temporal intervals. When a human participant compares two stimuli, the second auditory interval is compared against the decaying average of the first one. Raviv et al., suggested that errors in temporal discrimination arise during memory retrieval/decision making and not during memory encoding. A prediction of this model is that participants will have a better performance with a short ISI than with a long one.

Bayesian models offer a dynamical approach and take into consideration the representation of the two stimuli. The result of perceiving a duration, (the posterior distribution) is the product of the representation of the previous trial —and all the information collected about the underlying statistics until that point— (the prior distribution) and the current sensory input (the likelihood). Thus, in this model the prior is updated from trial to trial, whereas the posterior is modulated by the perception of both S and C. De Jong and colleagues [12] found that in comparing the duration of visual stimuli the influence of statistical context on time estimation is best explained by a Bayesian model using a Kalman filter, and thus discarded the IRM model. They found that the Type A effect is influenced by a dynamic prior that is sequentially updated by both stimuli: S and C.

In our own work, we showed that in discriminating two empty visual events with S < 200 ms, time distortions appear only if the ISI is shorter than about one second [22]. Here, we focused on the type A effect and tested how the factor that determine serial dynamics of relative event duration —relative position, relative distance in time and relative duration of S and C— contribute to generating temporal distortions. We used an S of 120 ms, and varied the ISI over four different intervals (400, 800, 1600, and 2000 ms).

Firstly, we swapped the order of presentation of S and C (Relative position factor). Secondly, we tested whether a long ISI increases the temporal accuracy (Relative distance factor). Finally, we tested whether the position of the longer stimulus modulates temporal accuracy (Relative duration factor), under the assumption that the contraction bias, which for short first stimuli should lead to a subjective expansion, applies independently of event type (S or C). We thus hypothesized that: 1) with an ever-changing C in first position, temporal accuracy and the magnitude of time distortions would increase, and participants would benefit to a markedly lesser extend from increased attention orienting for long ISIs, as the second event (S) would be already fully predictable. Hence, we expected an interaction between the two factors: stimulus presentation order and ISI; 2) if distortions in duration comparisons are mainly due the predictability features of S and C (trial-by-trial predictable vs. unpredictable), then the position of the longer stimulus should not modulate temporal perception.

Results verify hypothesis 1: the best Generalized Linear Mixed Model (GLMM) included an interaction between stimulus presentation order(<SC> or <CS>) and ISI. Increasing ISI reduces temporal distortions, more so for the <SC> group. Surprisingly, however, and contrary to our hypothesis 2, the relative position of the longer stimulus has important modulatory effects on temporal perception. Not all first-position events are subjectively expanded to the extent that they produce distortions in temporal judgment. Instead. time distortions tend to be generated when the first event in a series is shorter than the second event, independently of event type (S or C). Notably though, when the dynamic stimulus (C) is in first position, the ensuing distortion effect cannot be compensated by increasing ISI.

To dig deeper into the mechanics of TOEs and show the computational plausibility of the highlighted perceptual bias, we considered the fixed factors of the best GLMM and simulated our findings by implementing a Bayesian model using a Kalman filter [23, 24, 12]. Model results confirm the findings on human participants: first-position shorter events are disproportionately expanded. Our results contribute to clarifying the mechanics generating perceptual time distortions, by identifying a novel duration-dependent encoding inefficiency in human serial time perception.

## 2 Results

Two separate groups of human participants performed a 2IFC discrimination task comparing the duration of an S event against that of a C event (or vice versa), and deciding which stimulus was longer. To signal the onset and offset of each event, we used a short-duration blue disk (hence, S and C were empty visual stimuli, see Material and Methods section). For the <SC> experimental group, the S stimulus was displayed in the first position, and was shifted to the second position for the <CS> experimental group (Fig. 1a). The duration of the S event was kept constant (120 ms), whereas the duration of the C event varied, providing participants with three degrees of sensory evidence(weak ±Δ 20, medium ±Δ60, and strong ±Δ100 ms; Fig. 1b). We parametrically manipulated the ISI by using four durations: 400, 800, 1600, and 2000 ms.

**Figure 1:**
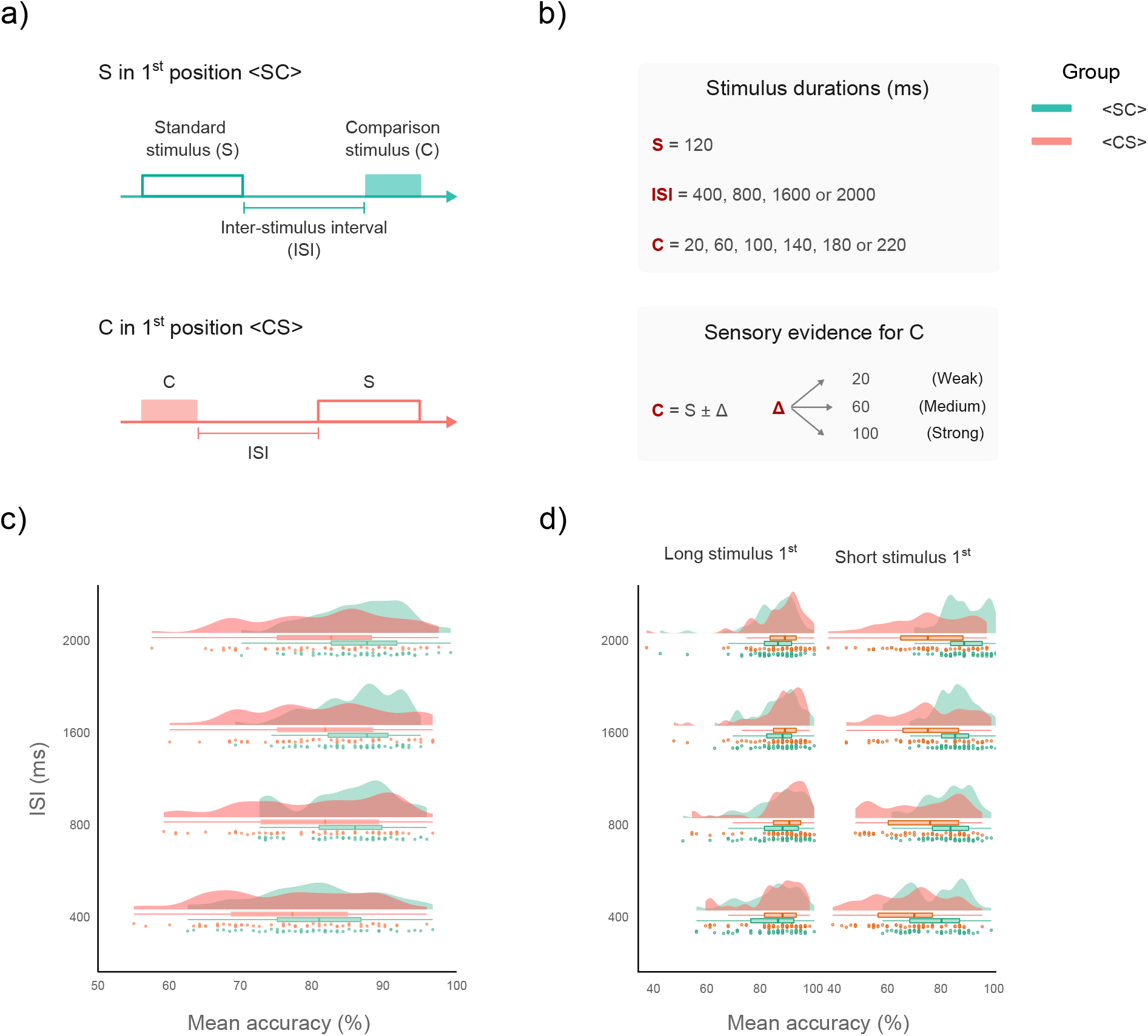
Two-interval forced choice (2IFC) task and accuracy. **a)** Timeline of events in the 2IFC task. For the <SC> group the Standard stimulus (S) was displayed in the 1^*st*^ position and in the 2^*nd*^ position for the <CS> group. **b)** The S stimulus had a fixed duration of 120 ms, whereas the Comparison stimulus varied trial-by-trail according to its level of sensory evidence: Weak, Medium, or Strong. We implemented four Inter-stimulus intervals (ISIs: 400, 800, 1600 and 2000 ms). **c)** Mean accuracy for each group and each ISI level. Data-points depict the mean accuracy for each participant. Box and density plots show the distribution of the mean accuracy for both <SC> and <CS> groups. The median is represented by the vertical line in the box plots, whereas the horizontal lines depict the interquartile range (IQR). Accuracy is higher for the <SC> group, however for both groups accuracy increased for ISI > 400 ms. **d)** Mean accuracy separated by the ordinal position of the longer stimulus. Accuracy for the <CS> group decreases when the 1^*st*^ stimulus is shorter than the 2^*nd*^ stimulus.

### 2.1 Temporal accuracy

Mean temporal accuracy was 84.39% (SD = 7.16) and 79.75% (SD = 9.79) for the <SC> and <CS> groups, respectively. Mean accuracy improved across the board for ISI conditions > 400 ms, however accuracy values were always lower for the <CS> than for the <SC> group (Fig 1c; Table 1). Raw accuracy was analyzed with GLMMs using a binomial parameter with a logit link function [25, 10]. We opted for a forward selection approach, and first used the magnitude of the difference between S and C as predictor, that is the Δ, which encodes three levels of sensory evidence, 20, 60, and 100 ms. We then added random intercepts for each participant. The model improved, indicating significant individual variability among participants (BF_01_ = 0). For the remaining models we always included random intercepts for each participant. To avoid convergence issues we refrained from adding more random effects.

**Table 1:**
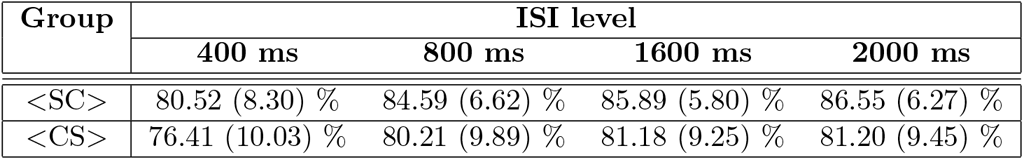
Mean accuracy. The table display the mean accuracy and the standard deviation (in parenthesis) for each group and each ISI level. For both groups, <SC> and <CS>, participants had their worst performance at the ISI_400_ condition. However, mean accuracy values are lower for the <CS> group than for the < SC> group.

| Group | ISI level |  |  |  |
| --- | --- | --- | --- | --- |
|  | 400 ms | 800 ms | 1600 ms | 2000 ms |
| <SC> | 80.52 (8.30) % | 84.59 (6.62) % | 85.89 (5.80) % | 86.55 (6.27) % |
| <CS> | 76.41 (10.03) % | 80.21 (9.89) % | 81.18 (9.25) % | 81.20 (9.45) % |

After, we added the ISI as an interaction with the Δ. The model again improved suggesting that the ISI factor modulates temporal accuracy differently depending on the amount of evidence for a difference between S and C: weak, medium, or strong (BF_01_ < 0.001). To assess whether the ordinal position of the longer stimulus modulates accuracy, we added this factor as an interaction. The model further improved, as the relative duration of first and second events, regardless of their experimental function (Standard or Comparison), significantly modulates temporal accuracy (BF_01_ < 0.001).

Finally, to test whether the order of S and C presentation has modulatory effects on accuracy (Fig 1d), we added Group as a fixed factor. The model improved significantly when the Group factor was added in interaction with the ISI factor, suggesting that temporal accuracy depends on the serial order of presentation of S and C (BF_01_ <0.001). Thus, the best model for explaining the raw accuracy included the interaction of the Δ, ISI, Group, and “Ordinal position of the longer stimulus” as predictors.

To examine the effects of relative stimulus duration, we deployed pairwise comparisons on the estimated marginal means (EMMs) of the best GLMM. Contrast analyses at each ISI level showed that for both groups <SC> and <CS> the ordinal position of the longer stimulus has modulatory effects on the accuracy (*p* = 0.0254; *p* < .0001; respectively). However, for the <SC> group post hoc analyses revealed statistically significant differences between EMMs only at the ISI_400_ level, whereas for the <CS> group we found statistically significant differences at each ISI level (all *ps* < .0001; Table 2), suggesting that an increasing ISI does not help suppressing distortions.

**Table 2:** Pairwise comparisons of estimated marginal means (EMMs) for each ISI level. The table display the results of the contrasts between the position of the longer stimulus conditions: 1^*st*^ and 2^*nd*^ position, at each ISI level. For the <SC> group the only statistically significant comparison was at the ISI_400_, whereas for the <SC> group all pairwise comparisons were statistically significant.

| Group | ISI | estimate | SE | df | z. ratio | p. value |
| --- | --- | --- | --- | --- | --- | --- |
| <SC> | 400 | 0.29 | 0.07 | Inf | 3.87 | < .0001 |
|  | 800 | 0.18 | 0.09 | Inf | 1.91 | < .0552 |
|  | 1600 | 0.07 | 0.10 | Inf | 0.72 | < .4677 |
|  | 2000 | -0.13 | 0.09 | Inf | -1.42 | < .1547 |
| <CS> | 400 | -2.41 | 0.20 | Inf | -11.74 | < .0001 |
|  | 800 | -2.45 | 0.21 | Inf | -11.55 | < .0001 |
|  | 1600 | -2.16 | 0.21 | Inf | -10.23 | < .0001 |
|  | 2000 | -1.89 | 0.21 | Inf | -9.01 | < .0001 |

Results of contrast analyses at the Δ levels showed that for the <CS> group all pairwise comparisons were statistically significant (all *ps* < .0001; Table 3). However, for the <SC> group results revealed a significant differences only at the Δ20 level (*p* < .0002; Table 3). These results suggest the existence of a perceptual bias that can be minimized by both an increase of the ISI or providing more sensory evidence (that is, Δ), but only if the first event has a constant duration.

**Table 3:** Pairwise comparisons of EMMs for each Delta level. The table display the results of the contrasts between the position of the longer stimulus conditions: 1^*st*^ and 2^*nd*^ position, at each Delta level. For the <SC> group the only statistically significant comparison was at the ISI_4_00, whereas for the <SC> group all pairwise comparisons were statistically significant.

| Group | Delta level | estimate | SE | df | z. ratio | p. value |
| --- | --- | --- | --- | --- | --- | --- |
| <SC> | 20 | 0.18 | 0.04 | Inf | 3.74 | < .0002 |
|  | 60 | 0.05 | 0.07 | Inf | 0.69 | < .4853 |
|  | 100 | 0.07 | 0.10 | Inf | 0.69 | < .4855 |
| <CS> | 20 | 1.54 | 0.05 | Inf | -11.74 | < .0001 |
|  | 60 | 1.04 | 0.06 | Inf | -11.55 | < .0001 |
|  | 100 | -0.65 | 0.09 | Inf | -7.01 | < .0001 |

### 2.2 Constant error (CE)

To obtain the temporal sensitivity and the magnitude of the time distortions (indexed via the PSE) of each group, we fitted GLMMs using the percentage of responses “C longer than S”. We used as predictors the fixed factors of the previous model, with the exception of the “ordinal position of the longer stimulus” factor. Here again, we used a binomial parameter. However, this time using a probit link function.

We first used the C stimulus as predictor, which encodes three degrees of sensory evidence: weak ±Δ 20, medium ±Δ60, and strong ±Δ100 ms. We then added random intercepts for each participant and each C stimuli (BF_01_ < 0.001; BF_01_ < 0.001, respectively). As the model improved, we included random intercepts for each participant. After this we added the ISI as a fixed factor. The model improved when we added an interaction between C stimulus and the ISI (BF_01_ < 0.001). Then, we added Group as a fixed factor (Fig. 2a). The model improved significantly when we added it as an in interaction with the ISI (BF_01_ < 0.001), suggesting that the probability of responding “C longer than S” depends on the ISI but also on the serial order of presentation of S and C. Thus, the best model included the interaction of the C stimulus, the ISI, and Group.

**Figure 2:**
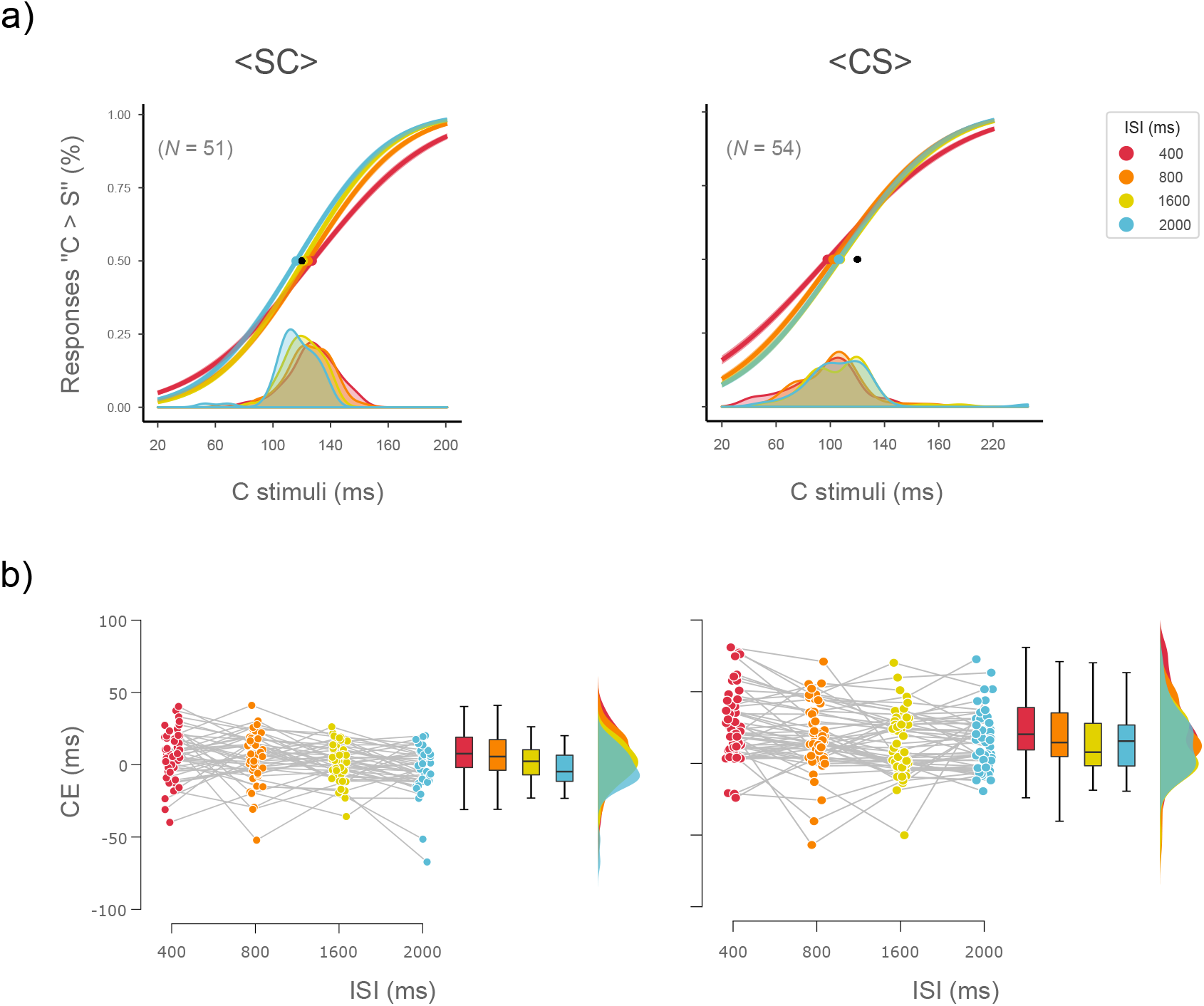
Constant error (CE) of human observers. **a)**Psychometric curves of human observers. Fitted curves modeling performance on each ISI level for both groups: <SC> and <CS>. The durations of the six C stimuli are plotted on the *x*-axis and the probability of responding “C longer than S” on the y-axis. Lines depict separate fits for each ISI condition, the black dot depicts the physical magnitude *ϕ_s_* of the Standard stimulus, the rest of the dots represent the point of subjective equality (PSE) for each ISI condition. Density plots show the subject-to-subject variability of the PSE for each ISI level. **b)** Data-points depict individual CEs for both <SC> and <CS> groups at each ISI level. Box-whisker, and density plots show the distribution of the CEs. The median is represented by the horizontal line in the box plots, whereas the bottom and top whiskers depict the IQR. a) For both groups the CE decreases with an increase of the ISI. However, CEs are larger for the(<CS>) than for the <SC> group.

To obtain the individual temporal sensitivity and the PSE, we fitted a GLM for each participant by using as predictors the fixed effects of the best GLMM. To obtain the exact magnitude of the time distortions we derived the CE from the PSE (see Material and Methods section). Results showed that, group-wise, CE values decrease with increasing ISI regardless of the group. However, CE values are higher for the <CS> than for the <SC> group (Fig. 2b, Table 4). Indeed, a repeated-measures Bayesian ANOVA revealed that best model for explaining these data was the model including the factors ISI and Group (BF_10_ = 5.7 ∗ 10^7^).

**Table 4:** Constant errors (CEs). The table display the mean CEs and the standard deviation (in parenthesis) for each group and each ISI level. For both groups, <SC> and <CS>, the CE decreased with an increase of the ISI. However, CEs were larger for the <CS> than for the <SC> group.

| Group | ISI level |  |  |  |
| --- | --- | --- | --- | --- |
|  | 400 ms | 800 ms | 1600 ms | 2000 ms |
| <SC> | 6.78 (17.45) | 4.52 (17.94) | 1.27 (13.11) | -4.23 (16.26) |
| <CS> | 25.46 (24.19) | 17.86 (23.46) | 13.45 (21.82) | 15.35 (20.24) |

### 2.3 Bayesian model

We implemented a Bayesian model to replicate our results by using a Kalman filter Petzschner & Glasauer, Glasauer & Shi, and de Jong et al., [23, 24, 12]. For each group we simulated data of 100 subjects using 120 trials at each ISI level (see Material and Methods section). We applied the best GLMM of the human observers to the simulated data. As with the human participants, we applied a GLM to each subject to obtain the individual temporal sensitivity and the PSE. To compare the responses of the human observers against the Bayesian observer’s responses, we obtained the root mean squared error (RMSEs). Results showed that the Bayesian observer’s responses successfully simulated the trend of results of the human observers: 1) the CEs decrease with an increase of the ISI; 2) CEs values are higher for the <CS> than for the <SC> group (Fig. 3a-b).

<SC> <CS>

**Figure 3:**
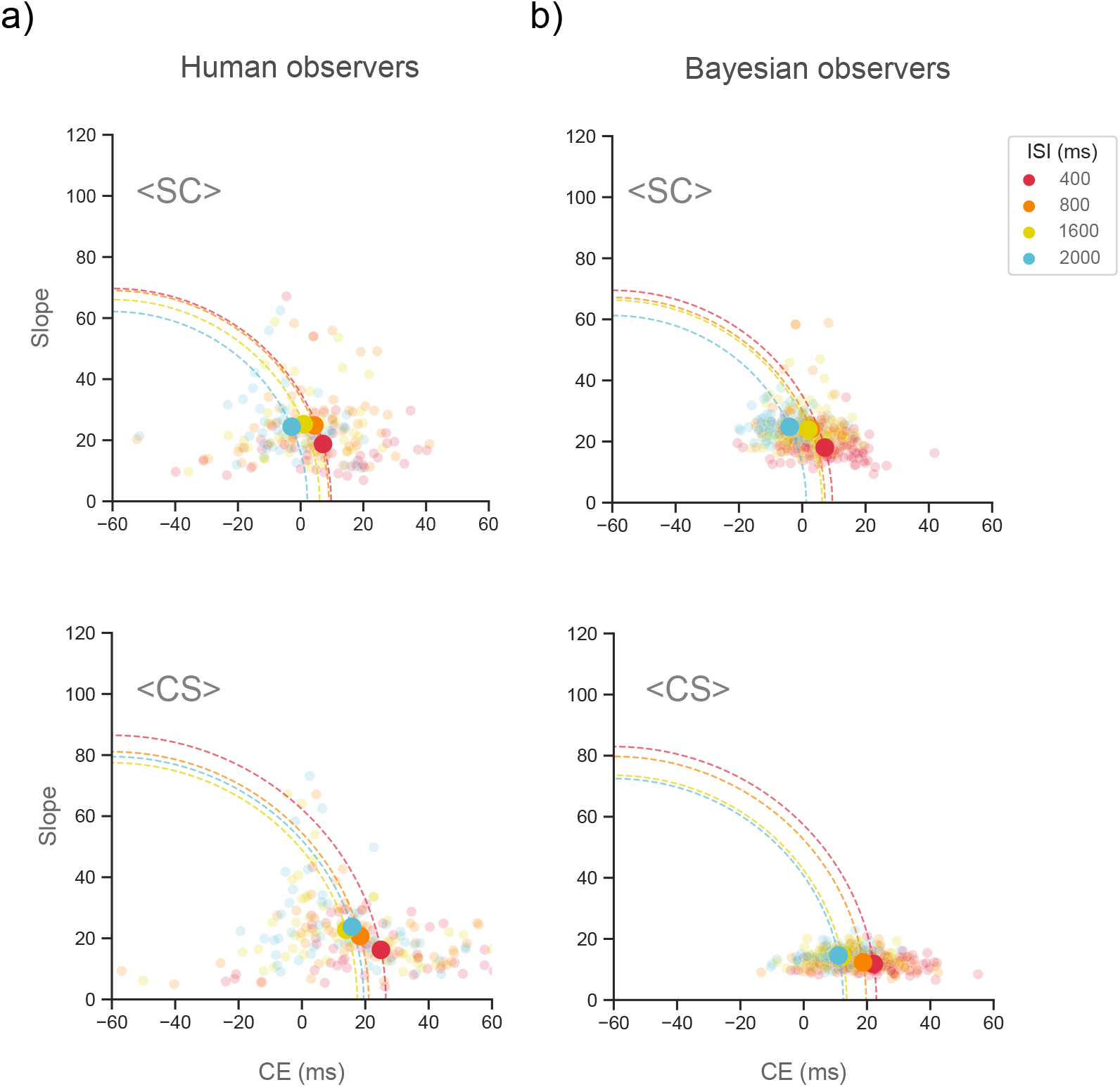
Root mean squared errors (RMSEs) of human and Bayesian observers. **a)** RMSEs of human observers. RMSEs are given by the distance from the origin and are depicted by a quarter circle. Any increase in the CE or the slope will lead to a larger radius. Big dots depict the intersection of the CE and the slope’s mean for each group and each ISI level. Small dots depict individual CEs and slopes values. CE mean values decrease as a function of the ISI regardless of the group (<SC> or <CS>). However, as we can see in the quarter circles, CE mean values are higher for the <CS> than for the <SC> group. **b)** RMSEs of Bayesian observers. Bayesian observer’s responses successfully simulated the main of results of the human observers: 1) the CEs decrease with an increase of the ISI, 2) CEs values are higher for the <CS> than for the <SC> group.

## 3 Discussion

The duration of an event can be distorted when the event is inserted in a series. Such effects, termed Time Order Error (TOE), constitutes one of the oldest and most investigated phenomena of subjective time perception [5, 26, 17]. Yet, the mechanics of TOE generation are still unknown. Since TOEs occur during serial discrimination tasks, we tested how event duration dynamics - relative position, distance in time and duration of two successive events - contributes to time distortions by flipping the positions of S and C events in two separate behavioral experiments.

We obtained three main findings. First, the interaction between stimulus order presentation and ISI verifies that, with an ever-changing and therefore unpredictable stimulus in first position, temporal accuracy decreases. These differences in accuracy lead to the Type A effect: short CEs for the <SC> group and large CEs for the <CS> group. Despite these differences, we replicated our finding that, by increasing the ISI between the first and second event, CEs decrease, although significantly less frequently for the <CS> group [22]. Dyjas et al., [18] found no significant statistical differences for the Type A effect between <SC> and <CS> presentation orders, as far as both visual and auditory modalities are concerned. Our findings contradict these results and show that at least when using empty visual events, the serial order of presentation <SC> and <CS> modulates temporal accuracy and CE.

Classical TOE studies suggest that the level of noise in the internal representation of the 1^*st*^ stimulus is larger than the 2^*nd*^ stimulus’ noise due to the encoding process and the maintenance of the 1^*st*^ stimulus in memory [27, 28]. Likewise, recent results in the auditory system propose a Bayesian model where the discrimination of two stimuli is done by comparing the second tone versus the decaying average of the first tone [14]. Thus, errors in temporal discrimination should be occur during memory retrieval and decision-making processes. Such a Bayesian model predicts that participants will have a better performance with a short ISI. However, our results show that increasing maintenance in memory does not have a detrimental effect on accuracy, on the contrary it has beneficial effects, possibly by reducing attentional blink effects. Indeed, our results are consistent with Grondin’s findings showing that the CE is reduced when the duration of the ISI is 1.5 seconds [9]. Grondin found that this benefit occurs when using both single and multiple visual standard stimuli. We suggest that these results can be explained by the beneficial effects of allocating attention in time, oriented to the encoding of unpredictable events [29, 30, 31]. Attention helps when S is in the first position, as it enhances the encoding of the C stimulus whose duration is unpredictable. We used this assumption to build our Bayesian model: allocating attention in time was modeled by decreasing the noise of the sensory input when the ISI increased.

The second finding concerns the preeminence of the ordinal position of the longer stimulus —independently of event type (S or C)— in driving accuracy. This prefigures a novel serial order bias in serial perception based on duration-dependent relative positions of stimuli. When the first stimulus in a series is shorter than the second stimulus, regardless of whether it is S or C, participants were biased to say that the first event was longer, consistently making mistakes. We found this effect in all experimental conditions of the <CS> group, but for the <SC> group it was only present at the shortest ISI (400 ms). We also showed that the serial order bias or perceptual glitch arises at all sensory levels of the <CS> group but only at the weak sensory level of the <SC>. These patterns of results explain why the magnitude of the time distortions (indexed via the CE) are larger for the <CS> than for the <SC> group. The serial order bias can be minimized by an increase of the ISI or an increase in the level of the sensory evidence if the first stimulus is predictable (<SC> group). However, when a dynamic stimulus is displayed in first position (<CS> group) such a bias is at ceiling and leading to temporal errors that will increase the magnitude of the time distortions across the board.

Third, our Bayesian model successfully simulated our human behavioral findings and showed that the Type A effect arises under sensory uncertainty because of the highlighted serial perceptual encoding inefficiency. Our modelling captures the idea that the temporal stimulus’s perception in the visual system is shaped not only by sensory noise but also by a perceptual bias that systematically makes participants expand the first stimulus in a series if unpredictable. Naturally, if such stimulus is already longer than the second, the bias would not be visible as it would not lead to perceptual mistakes. By simulating our behavioral results with a Bayesian model, we provide more evidence to show that time estimation and duration discrimination are a dynamic process that not only take in consideration the current sensory information of the two stimuli but also the information of previous trials. Our findings align with recent results showing that the type A effect is best modeled with a Bayesian model using a Kalman filter [12].

The replication of our behavioral results with a Bayesian model offers an insight on the computations that the human brain might use as a strategy for temporal discrimination. At the same time, it shows how this computation is affected by sensory noise, the allocation of attention in time, and a hitherto unknown perceptual bias. On a final note, our behavioral results and simulations highlight the importance of the ISI on a 2IFC paradigm. De Jong et al., [12] and Raviv et al., [14], proposed powerful models for temporal discrimination on the visual and the auditory system, respectively, but implicit in their experimental paradigms and models is the employment of a long ISI. De Jong and colleagues implemented and ISI of 1000 ms, whereas Raviv et al., used and ISI of 950 ms. Here, in order to take in consideration, the effects of ISI, we modeled the ISI’s effects by decreasing the noise of the sensory input when the ISI increased. To do that, we implemented different distributions for the internal representation’s noise.

As our stimuli used visual intervals at the bottom of the sub-second scale (120 ms), the question remains as to whether the perceptual serial order glitch would disappear for stimulus intervals in the supra-second range, and also whether it is present in other sensory modalities, besides vision. Future research is needed to uncover the physiological basis of such a strong, implicit expectation about the temporal statistics of incoming stimuli which can drive humans to distort time perception under uncertainty.

## 4 Materials and Methods

### 4.1 Participants

The experiment was organized as a between-subject design, with separate groups for the position of the stimuli: S in first position (<SC>) and C in first position (<CS>). Part of the results of the <SC> group were previously published [22]. This data-set has a sample of 52 participants (34 female; ages: 18-33; mean age: 24.42). One participant was removed due to chance level accuracy (< 55%). Therefore, the final sample included the data from 51 participants (33 female; ages: 18-33; mean age: 24.45). For the <CS> group we had an initial sample of 58 participants (45 female; ages: 18-37; mean age: 25.41). Four participants were removed due to chance level accuracy (< 55%). Therefore, the final analysis included the data from 54 participants (41 female; ages: 18-33; mean age: 25.31). In total, we report on the behavior of 105 participants. For the analyses of the slope and constant error (CE), participants were excluded when one of the dependent variables had a value with three standard deviations above (or below) the mean. Thus, for the analyses of the slope and CE, nine participants were excluded following this procedure.

Individuals were recruited through online advertisements. Participants self-reported normal or corrected vision and had no history of neurological disorders. Up to three participants were tested simultaneously at computer workstations with identical configurations. They received 10 euros per hour for their participation.

### 4.2 Design

We used a classical interval discrimination task by implementing a 2IFC design, where participants were presented with two visual durations: S and C [6, 32]. S had a magnitude of 120 ms. For the <SC> group 5 was always displayed in the first position, but it was shifted to second position for the <CS> group (Fig 1a). In both groups, we used three magnitudes for the step comparisons Δ between S and C: 20, 60, and 100 ms. We derived the magnitudes for the C stimuli as S ± Δ, which resulted in the next C intervals: 20, 60, 100, 140, 180, and 220 ms. C stimuli were randomized on a trial-by-trial base.

We used the same four ISIs for both groups: 400, 800, 1600, and 2000 ms. For each trial, the inter-trial interval (ITI) was randomly chosen from a uniform distribution between 1 and 3 seconds. Participants judged whether the S or C stimulus was the longer duration. They responded by pressing one of two buttons on an RB-740 Cedrus Response Pad (http://cedrus.com) and were provided with immediate feedback on each trial.

### 4.3 Stimuli and Apparatus

Stimulus duration was determined as a succession of two blue disks with a diameter of 1.5° presented on a gray screen [33]. Empty stimuli were implemented to ensure that participants were focused on the stimuli’s temporal properties [34]. All stimuli were created in MATLAB R2018b (http://mathworks.com), using the Psychophysics Toolbox extensions [35, 36, 37]. Stimuli were displayed on an ASUS monitor (model: VG248QE; resolution: 1,920 x 1,080; refresh rate: 144 Hz; size: 24 in) at a viewing distance of 60 cm.

### 4.4 Protocol (Task)

The experiment was run in a single session of 70 minutes. Participants completed a practice set of four blocks (18 trials in each block). All sessions consisted of the presentation of one block for each ISI condition. Each block was composed of 120 trials. For each ISI the C intervals were presented in random order. Each ISI block was also presented in random order, each randomization was unique.

To avoid fatigue, participants always had a break after 60 trials. Each trial began with a black fixation cross (diameter: 0.1°) displayed in the center of a gray screen. Its duration was randomly selected from a distribution between 400 and 800 ms. After a blank interval of 500 ms, S was displayed and followed by an ISI. After this, C was displayed. Participants were instructed to compare the interval of the two stimuli by pressing the key “left”, if S was perceived to have lasted longer, and the key “right” if C was perceived to have lasted longer. After responding, they were provided with immediate feedback: the fixation cross color changed to green when the response was correct, and to red when the response was incorrect.

### 4.5 Data analysis

Data cleansing was implemented with Python 3.7 (http://python.org) using the ecosystem SciPy (http://scipy.org). GLMMs were fitted in R [38] using the *lme4* (package version 1.1.21). Raincloud plots were created using the *raincloud* function for R [39]. All data analyses and simulations, whether using Python, were performed in Jupyter Lab (http://jupyter.org). The annotated notebooks can be consulted at Open Sciecne Framework (OSF) (https://osf.io/qnj3t/).

#### 4.5.1 General Linear Mixed Models (GLMMs)

We modeled our behavioral data with GLMMs to estimate a single model across all subjects, and distinguish within- and between-subjects errors [40, 25, 10]. To fit the GLMMs we input the responses as a whole [10, 12]. Raw accuracy was analyzed with GLMMs using a binomial parameter with a logit link function. To compute the temporal sensitivity and the PSE we fitted GLMMs using as dependent variable the percentage of responses “C > S”. We used again a binomial parameter but this time using a probit function.We calculated the expected value of the responses as follow:

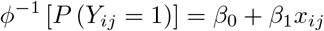

Where *X_ij_* is the Comparison stimulus’ duration, *Y_ij_* the response variable for subject *i* and trial *j*. If the C stimulus is judge longer than S, then *Y_ij_* = 1; but *Y_ij_* = 0, if C is shorter than S. The probability of the response “C longer than S” *P*(*Y_ij_*) = 1 is linked with the linear predictor via the probit link function *ϕ*^−1^. The fixed-effect parameters *β*_0_ and *β*_1_ are the intercept and the slope, respectively. The *β*_1_ is an index of the temporal precision, which is also called the Just Noticeable-Difference (JND) [10]. The PSE is a function of both parameters:

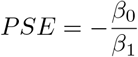

We derived the CE as the difference between the PSE and the magnitude *ϕ_s_* of the Standard duration: CE = PSE —*ϕ_s_*, and CE = *ϕ_s_* − PSE, for the <SC> and <CS> groups, respectively.

To apply Model Comparison (BMC) to the GLMMs and decide between models, we compute the Bayes factors using the Bayesian Information Criterion (BIC) [41]:

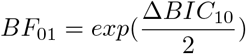

#### 4.5.2 Bayesian modelling

We implemented a Bayesian model to replicate our results by using a Kalman filter. We based our Bayesian model on the work of Petzschner & Glasauer, Glasauer & Shi, and de Jong et al., [23, 24, 12]. In this model the prior represents the intervals stored in memory (that is, the internal representation of previous trials), the likelihood is the current sensory input, and the posterior is the current estimate or *percept*. To run our Bayesian model and use it for temporal discrimination, we used the duration of both stimuli —S and C— as inputs for this model. To do that we used the representation of the stimulus’ duration on a logarithmic scale and added some Gaussian noise:

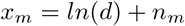

Where d is the physical duration of the stimulus (S or C) in a linear scale and *x_m_* is the internal noisy representation. The random variable *n_m_* represents the normally distributed measurement noise 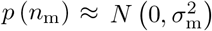 [42]. To run simulations for individual participants we randomly selected values for 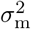 from a truncated normal distribution. Note that the magnitude of 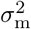 is the temporal sensitivity of each participant. The model compared on a trial basis the logarithmic representations of S and C, and yielded 0 or 1:

- if “S > C”, the model yielded 0
- if “C > S”, the model yielded 1

when a new duration —indexed by n— is perceived it is represented by the likelihood function, which is a Gaussian distribution with 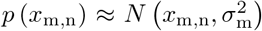. Note that S and C are the mean of the priors that emerged from perceiving both stimuli. The prior too is modelled as a Gaussian distribution: 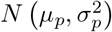. To estimate a stimulus’ duration the prior is updated through a weighted average of the previous prior distribution and the currently sensed likelihood. For each measurement the update step is modeled by the formulation of the Kalman filter for a 1D first-order system:

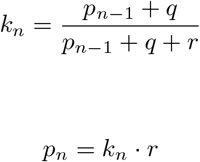

Where *r* is the is the uncertainty of the current likehood 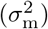 and *p*_*n*−1_ the uncertainty of the previous prior 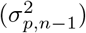. Thus, the Kalman gain (*k*) of a new observation is determined by both uncertainties and a process variance *q*. The variance of the prior system *p_n_* is updated by the product of the Kalman gain and r, whereas the prior mean *μ_p,n_* is updated as follow:

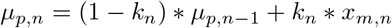

To simulate our behavioral results, we used three assumptions for modelling the noise *n_m_* associated with the internal representation *x_m·_* 1) We assumed that *n_m_* decreases as the duration of the ISI increases. That is, the noise *n_m_* associated with the second stimulus was larger for the ISI_400_ than for the ISI_2000_ condition. 2) We assumed that independently of the group (<SC> or <CS>) or even type (S or C) the stimulus’s noise come from two different distributions. Thus, we used a distribution for the “Longer stimulus in 1^*st*^ position” stimuli’ noise and a second distribution for the “longer stimulus in in 2^*nd*^ position” stimuli’ noise. 3) Because the perceptual bias is stronger at the weak sensory level than the other two sensory levels, we used a truncated normal distribution for *n_m_* at the weak sensory level (Δ20). For the <SC> group this bias affected the (+Δ20) trails, but for the group <CS> group affected the the (-Δ20) trials. See the annotated notebooks at OSF (https://osf.io/qnj3t/).

For each group we generated data for 100 participants with 120 trials for each ISI level. For both groups we kept a constant value for *q* (q = 1.5).

#### 4.5.3 Root Mean Squared Error (RMSE)

To compare the performance of human participants against the Bayesian obverver’s responses, we computed the RMSE which is given by the standard deviation (SD) and the bias: *RMSE*^2^ = *SD*^2^ + *bias*^2^, where the SD is the slope and the bias is the CE [19, 43]. As the RMSE can be written as the standard equation of the circle, it provides an effective geometric and graphical description to track changes on the CE and explore the trade-off between the CE and the temporal precision. RMSEs are given by the distance of the origin, and are depicted by a quarter circle. Because we had negative values for the CE, we took −60 as the origin point instead of 0. Thus, to find the circle’s intercept on the x-axis, that is the axis of the CE, we took the absolute distance between −60 and the CE.

## Data availability

Anonymized data are available at the Open Science Framework (https://osf.io/qnj3t/).

